# Local activation of focal adhesion kinase orchestrates the positioning of presynaptic scaffold proteins and Ca^2+^ channel function to control glucose dependent insulin secretion

**DOI:** 10.1101/2021.12.21.473760

**Authors:** Nicole Hallahan, Kylie Deng, Dillon Jevon, Krish Kumar, Jason Tong, Wan Jun Gan, Clara Tran, Marcela Bilek, Peter Thorn

**Affiliations:** Charles Perkins Centre, School of Medical Sciences, University of Sydney, Camperdown, 2006, Australia; Mechanobiology Institute, National University of Singapore, Singapore; School of Physics, University of Sydney, Camperdown, 2006, Australia; School of Aerospace, Mechanical and Mechatronic Engineering, University of Sydney, 2006, Australia; Sydney Nanoscience Institute, University of Sydney, Camperdown, 2006, Australia

## Abstract

A developing understanding suggests that spatial compartmentalisation of components the glucose stimulus-secretion pathway in pancreatic β cells are critical in controlling insulin secretion. To investigate the mechanisms, we have developed live-cell sub-cellular imaging methods using the organotypic pancreatic slice. We demonstrate that the organotypic pancreatic slice, when compared with isolated islets, preserves intact β cell structure, and enhances glucose dependent Ca^2+^ responses and insulin secretion. Using the slice technique, we have discovered the essential role of local activation of integrins and the downstream component, focal adhesion kinase, in regulating β cells. Integrins and focal adhesion kinase are exclusively activated at the β cell capillary interface and using *in situ* and *in vitro* models we show their activation both positions presynaptic scaffold proteins, like ELKS and liprin, and regulates glucose dependent Ca^2+^ responses and insulin secretion. We conclude that focal adhesion kinase orchestrates the final steps of glucose dependent insulin secretion within the restricted domain where β cells contact the islet capillaries.

## Introduction

The intrinsic stimulus secretion coupling cascade in pancreatic β cells is well understood through extensive *in vitro* experimentation (Rorsman and Ashcroft, 2018). However, within the native islets of Langerhans numerous external factors intersect with this signal cascade to further control secretion (Lammert and Thorn, 2020; Meda, 2013). The impact of some factors, such as gap junctions between endocrine cells, are well understood (Benninger et al., 2011). Less well understood is the impact of the islet microenvironment on β cell structural organisation and function (Lammert and Thorn, 2020) and how this intersects with the known stimulus secretion pathways.

Accumulating evidence suggests that the region where β cells contact the islet capillaries is specialised for secretion (Gan et al., 2017; Low et al., 2014). β cells, within intact islets, make a discrete point of contact with the extracellular matrix that surrounds the capillaries. This point of contact is the target for insulin granule fusion (Low et al., 2014) and is enriched in presynaptic scaffold proteins, like liprin and ELKS and therefore has characteristics analogous to a neuronal presynaptic domain (Lammert and Thorn, 2020; Low et al., 2014; Ohara-Imaizumi et al., 2019; Ohara-Imaizumi et al., 2005). Recapitulating this domain by culture of β cells on extracellular matrix coated dishes shows that local activation of integrins are the target for insulin granule fusion (Gan et al., 2018) and local control of microtubules regulates these secretory hot spots (Trogden et al., 2021). Although the mechanisms are not known this work suggests that presynaptic scaffold proteins, and perhaps microtubules, control granule targeting to this capillary interface.

Just like neurotransmitter release, Ca^2+^ is the dominant regulator of insulin secretion principally by Ca^2+^ entry through voltage sensitive Ca^2+^ channels (Schulla et al., 2003). We know from other systems that the location of Ca^2+^ channels relative to sites of granule fusion are critical to stimulus secretion coupling (Nanou and Catterall, 2018; Stanley, 1997). Ca^2+^ channels are typically regulated by intracellular Ca^2+^ concentrations leading to positive and negative feedback to control channel opening (Zühlke et al., 1999). These actions control the amplitude and temporal kinetics of local subcellular Ca^2+^ concentrations which in turn regulate exocytosis (Nanou and Catterall, 2018). In neurones, a presynaptic scaffold protein complex tethers synaptic vesicles and collocates Ca^2+^ channels to the presynaptic domain (Sudhof, 2012); whether similar mechanisms exist at the capillary interface of β cells is unknown.

In β cells there is functional, *in vitro*, evidence for close association of Cav1.2, Ca^2+^ channels and insulin granule exocytosis (Bokvist et al., 1995; Gandasi et al., 2017; Pertusa et al., 1999) and structural evidence for protein links between the Cav1.2 channels and syntaxin 1A (Wiser et al., 1999); a key SNARE protein required for granule fusion. This evidence is based on single, isolated β cells where capillary contacts are not present and the normal environmental cues of the islets are lost. Immunostaining β cells in the more intact environment of pancreatic slices shows that syntaxin 1A (Low et al., 2014) has an even distribution across the β cell plasma membrane and no enrichment at the capillary interface. This evidence therefore discounts a simple model where insulin secretion is regulated by colocalisation of syntaxin 1A and Cav1.2 at the capillary interface. Instead, there is recent evidence that the scaffold protein ELKS interacts with the β subunit of the Ca^2+^ channel (Ohara-Imaizumi et al., 2019). Furthermore, although the work was carried out in isolated islets, which lack capillaries, there was evidence that the coupling between ELKS and the β subunit enhanced the Ca^2+^ response at residual capillary structures (Ohara-Imaizumi et al., 2019), consistent with the idea that localised synaptic-like regulation of Ca^2+^ and exocytosis might exist in β cells. However, the mechanisms that organise and control the positioning of these presynaptic scaffold proteins is unknown.

The emerging picture therefore is that spatial compartmentalisation is a key attribute of stimulus-secretion coupling in pancreatic β cells. The capillary interface of β cells is a region enriched in presynaptic scaffold proteins, is the target for insulin granule fusion and might be a region where Ca^2+^ channels are regulated. However, progress in this area is hampered by the difficulties in imaging single β cells within the islet environment.

To this end, the pancreatic slice is as an important advance with a closer preservation to native islet structure than isolated islets (Gan et al., 2017; Meneghel-Rozzo et al., 2004). Analogous to organotypic brain slices, pancreatic slices maintain complex cell-to-cell arrangements that are likely to be important for overall islet control such as an intact islet capillary bed (Cohrs et al., 2017; Gan et al., 2017). In addition, the local microenvironment around each endocrine cell is maintained, with each cell contacting the capillary and other endocrine cells. This promotes a distinct subcellular polarisation in β cells that is likely to impact on cell function (Gan et al., 2017) with recent evidence the same organisation is present in rodent and human islets (Cottle et al., 2021). To date the pancreatic slice has been used in fixed-cell studies (eg (Gan et al., 2017) and functional studies, for example looking at coordination of Ca^2+^ responses in β cells across the islet (Stožer et al., 2013). In principle, the slice is the ideal platform for live-cell sub-cellular studies of the effects of β cell organisation on glucose dependent responses. However, preservation of function in slices has proved difficult and to date single cell, live-cell work (eg (Low et al., 2014; Ohara-Imaizumi et al., 2019)) have relied on isolated islets where capillaries are damaged and fragmented (Irving-Rodgers et al., 2014; Lukinius et al., 1995).

Here, we have developed the pancreatic slice preparation for live-cell sub-cellular imaging of β cell responses to glucose. Compared to isolated islets we show; slices demonstrate local activation of integrins and focal adhesions at the capillary interface of β cells, preserve enrichment of presynaptic scaffold proteins and have highly targeted insulin granule fusion to the capillary interface, fast Ca^2+^ spikes at low glucose concentrations and fast Ca^2+^ kinetics in response to glucose elevation with very fast intracellular Ca^2+^ waves that originate at the capillary interface.

These distinct responses in pancreatic slices demonstrate the importance of β cell organisation and the crucial role of the capillary interface in glucose dependent responses with evidence this is likely to be initiated by activation of the integrin/focal adhesion pathway. To test this mechanism, we used a range of interventions to block integrins and focal adhesion kinase (FAK) all of which consistently inhibit glucose dependent Ca^2+^ responses and insulin secretion. Furthermore, the same interventions disrupt the positioning of the presynaptic scaffold proteins ELKS and liprin. Importantly we show high potassium secretion and Ca^2+^ responses are not affected demonstrating the integrin/FAK pathway is a key and selective mediator of glucose dependence.

Together our data demonstrates that FAK is a master regulator of glucose induced insulin secretion that controls the positioning of presynaptic scaffold proteins and the functioning of Ca^2+^ channels.

## Results

The native environment of the islets of Langerhans is recognised as a key factor in controlling β cell behaviour (Lammert and Thorn, 2020) and accumulating evidence demonstrates that organotypic pancreatic slices preserve much of the normal structure of islets (Cottle et al., 2021; Meneghel-Rozzo et al., 2004) making them a preferred platform to drive new insights into the control of β cells and insulin secretion. The striking characteristic of islets within slices is the preservation of the rich capillary bed (Gan et al., 2017) which, contrasts with the loss of endothelial cells and capillaries in the more usual method of enzymatic islet isolation (Lukinius et al., 1995).

Using an immunostaining approach, as expected, because endothelial cells are the only source of intra-islet extracellular matrix (ECM) (Nikolova et al., 2006), the distribution of ECM was markedly different between pancreatic slices and isolated islets (Fig 1). In the isolated islets laminin was not associated with any specific structure but instead was dotted throughout the islets (Fig 1A). In pancreatic slices the ECM, identified by laminin, was strongly enriched around the islet capillaries and in the islet capsule (Fig 1B). Consistent with this disruption we observed a reduction in laminin area and a reduction in CD31 (endothelial cell marker) immunostaining (Fig1C).

**Figure 1.**
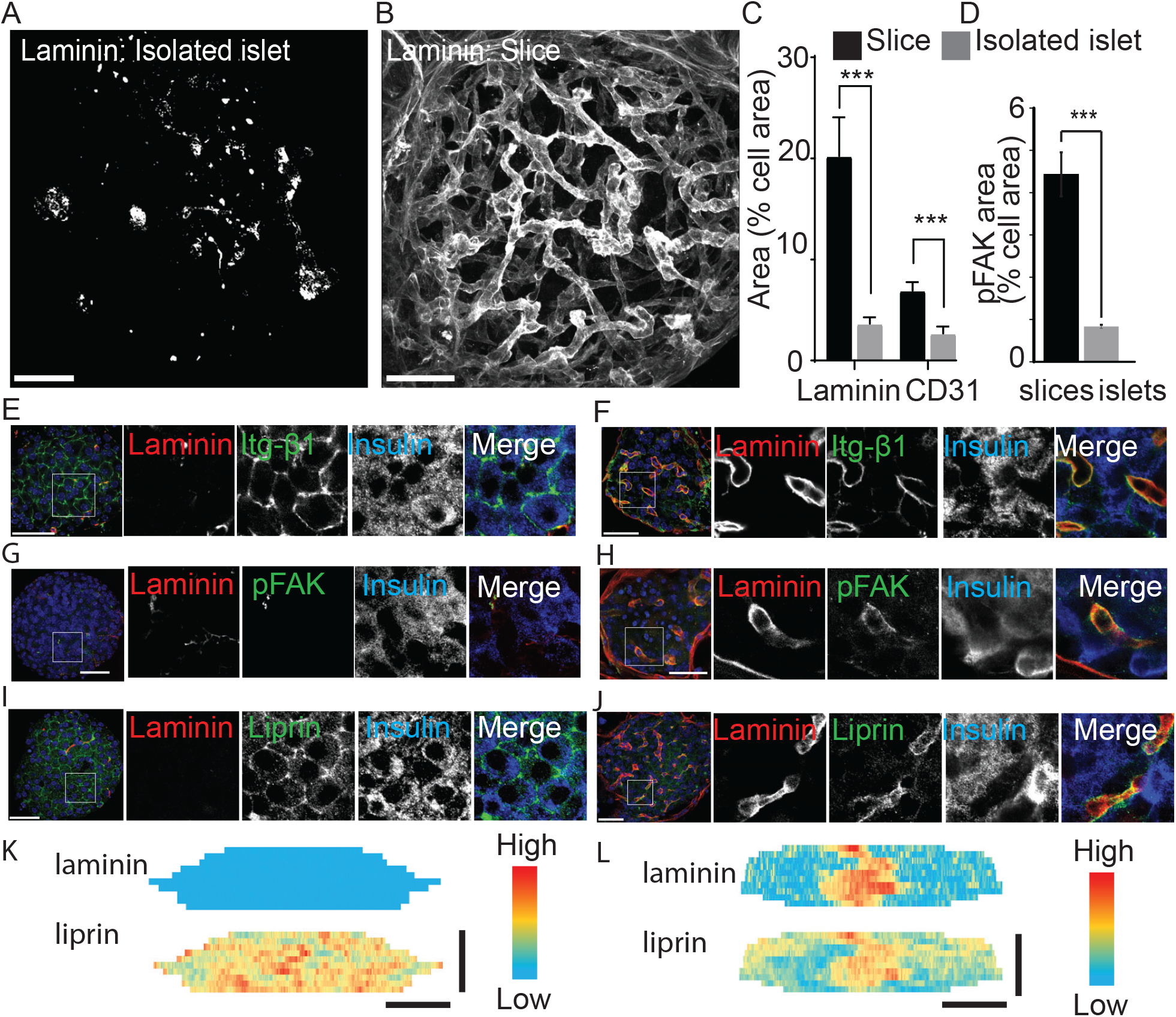
Pancreatic slices have an intact capillary bed. Integrin β1, phosphorylated FAK and liprin are enriched at the β cell capillary interface. Pancreas slices and isolated islets were cultured for 24 hours prior to fixation and immunostaining. Representative 3D projection of the ECM protein laminin through (A) an isolated islet and (B) an islet embedded in a 200 μm thick pancreas slice. (C) quantification of laminin and CD31 immunofluorescence, normalised to cell area (insulin + DAPI) in the corresponding Z-planes showed a significant loss of both proteins in isolated islets compared to slices. (n=3 and 2-3 islets analysed per mouse, mean ± SEM, student’s t-test p<0.001). Scales 40 µm. (D) phospho-FAK immunostaining shows that active focal adhesions are significantly reduced in area (compared to total cell area, insulin + DAPI) in β cells in isolated islets (E) (n=29 cells in slices 112 cells in islets, Student’s t-test p<0.01). Scales 40 µm. (E,G,I) immunostaining in islets for integrin β1, laminin and liprin shows lack of organisation compared to the enrichment at the capillary interface of β cells in slices (F,H,J). Heatmaps of the total surface area of single β cells show the focussed enrichment of liprin-α co-incident with laminin staining in (L) slices that is lost in (K) isolated islets.

ECM activates integrin-mediated responses in β cells (Gan et al., 2018; Parnaud et al., 2006; Rondas et al., 2012), we therefore sought to define the subcellular responses in β cells in the two preparations. We observed tight alignment of integrin β1 with laminin-stained capillaries in slices (Fig 1F) and consistent with the relative loss of ECM proteins in isolated islets (Fig 1A) we observed a misdistribution of integrin β1 (Fig 1E), although interestingly, in isolated islets, integrin β1 was still present but was now all around the cells.

Phosphorylated FAK, (phospho-FAK) provides a read out of focal adhesion activity (Rondas et al., 2012). Immunostaining in the slices showed a local enrichment of phospho-FAK at the capillary interface, consistent with the localisation of capillary ECM (Fig 1H). In isolated islets phospho-FAK is also enriched at the residual capillaries (Fig 1G) but, using an area analysis, we observe a significant and approximately 5-fold decrease in area occupied by phospho-FAK in the islets compared to slices (Fig 1D). This data shows that the disruption in ECM in the isolated islets does affect the function of β cells, in this case, reducing focal adhesion activity, as measured by phosphorylation. We also show that β cell structure is affected, lateral β cell contacts were still maintained, as shown by E-cadherin immunostaining but Par3, normally located in the apical region of β cells, away from the capillaries (Supplemental Fig 1) showed diffuse non-polar organisation in isolated islets (Supplemental Fig 1).

A developing understanding in this area is that the capillary interface of pancreatic β cells has similarities to the presynaptic domain of neurons including the enrichment of synaptic scaffold proteins, like liprin and ELKS (Gan et al., 2017; Lammert and Thorn, 2020; Low et al., 2014; Ohara-Imaizumi et al., 2005). *In vitro*, we have previously shown that local integrin activation is a primary factor in causing the clustering of liprin (Gan et al., 2018). Using immunostaining in pancreatic slices, as expected, liprin showed enrichment at the capillary interface (stained with laminin) and very little staining in other regions around the β cell (Fig 1J, L). In contrast, in isolated islets, liprin was dispersed and located all around the β cell surface (Fig 1I, K).

We conclude that the organotypic slice preserves the secondary structure of the islet, such as the capillary bed and the polarised structure of the β cells. Functionally this translates into a local positioning of integrins and the local activation of phospho-FAK, both of which are significantly disrupted in isolated islets. We therefore set out to use the slice as a platform to test the functional consequences of preserved β cell structure and activation of the integrin/FAK pathway.

### Organotypic slices have significantly enhanced glucose sensitive insulin secretion

Our past work with isolated islets demonstrated that insulin granule fusion is targeted to the interface of the remnant capillaries in isolated islets (Low et al., 2014). The method uses a dye-tracing technique to identify in space and time the fusion of individual secretory granules induced by a step increase in glucose from 2.8 to 16.7 mM. Here we repeat those findings and record each fusion event over a 20-minute stimulus with high glucose (Fig 2A) but now, in parallel experiments, compare the distribution of events obtained using pancreatic slices (Fig 2B). In both preparations targeting to the capillary interface of β cells is observed but in the slice preparation the targeting is significantly enhanced to the extent that nearly 80% of all granule fusion events occur in this region (Fig 2C, D). The greater precision in targeting of insulin granule fusion is consistent with the tight focus of phospho-FAK enrichment in slices (Fig 1H) and with previous *in vitro* reports that integrin activation drives granule targeting (Gan et al., 2018).

**Figure 2.**
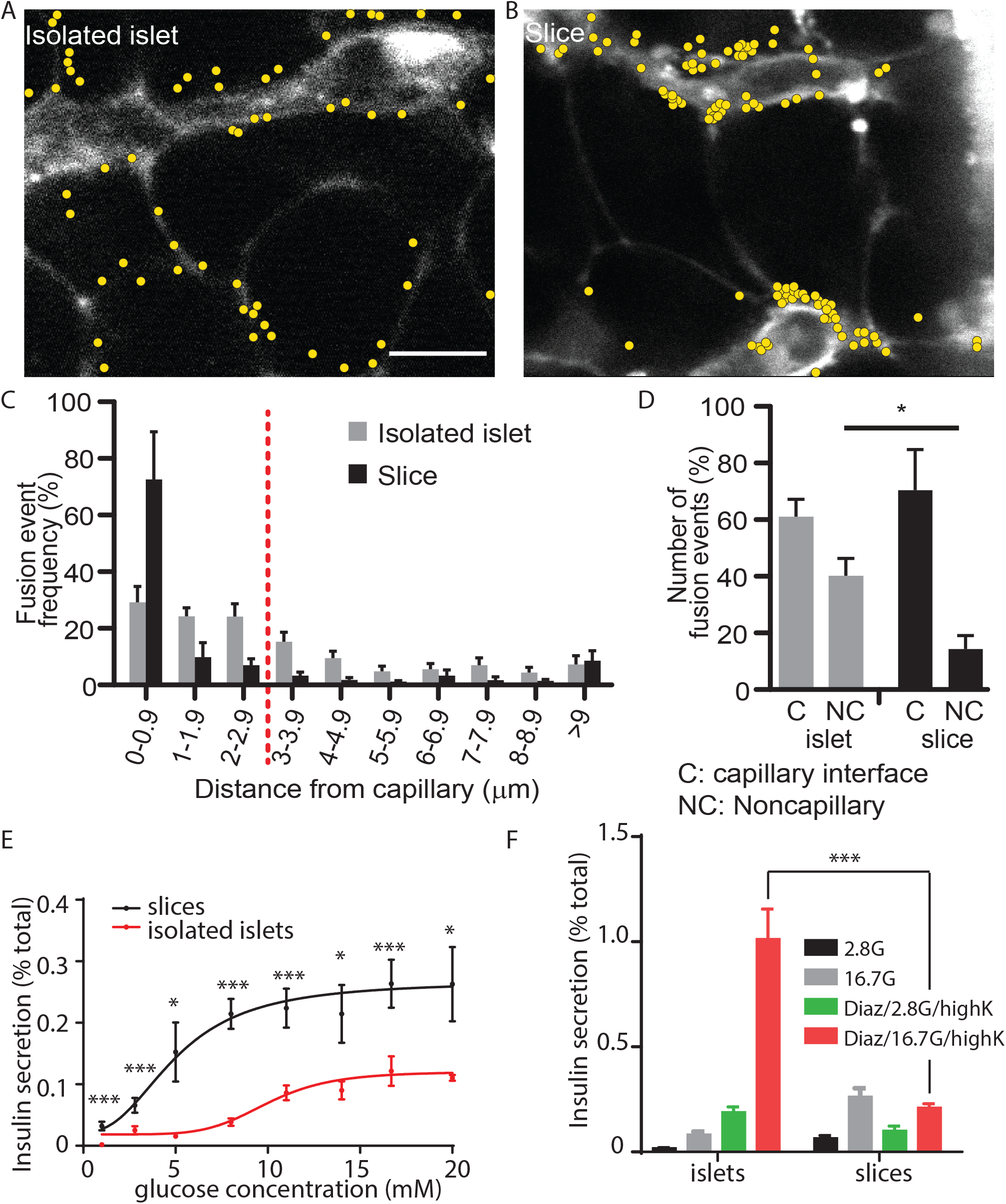
Glucose-stimulated insulin granule fusion and secretion in isolated islets and pancreas slices. (A) Isolated islets and (B) pancreas slices, bathed in an extracellular dye (sulforhodamine B, SRB), were stimulated with 16.7 mM glucose to induce insulin granule fusion, which is recorded as the sudden and transient appearance of bright spots of fluorescence (Low et al 2014). Continuous recording of two-photon images over 20 min of glucose stimulation led to many exocytic events, which were identified and marked on the images with yellow dots. (C) Slices (n=6 slices) had a strong bias of fusion events towards the vasculature, while fusion events in isolated islets (n=6 islets) were more spread out. (D) Fusion events in isolated islets and slices were classified as either occurring at the capillary face (<2.9 μm, C) or elsewhere on the cell membrane (>2.9 μm, NC). All data are represented as the mean ± SEM (n=3), significance determined by Student’s t tests, p< 0.05. (E) Dose-dependent glucose-stimulated insulin secretion normalised to total cellular insulin content shows that isolated islets are less sensitive to low glucose concentration and secrete a lower proportion of their total content compared to islets in pancreas slices (n=3-14 mice at each point, Student t test comparing responses at each glucose concentration were significant p<0.05). The lines are non-linear best fit dose-response curves with a fitted EC_50_ of 10.2 mM for islets and 8.6 mM for slices. (F) Islets and slices were incubated either with glucose alone at 2.8mM or 16.7mM glucose or in the presence of 250 µM diazoxide where secretion was subsequently stimulated by raising extracellular potassium. The response in the presence of diazoxide and 16.7 mM glucose, which reflects glucose amplification was significantly greater in islets compared to slices (n=6-13 islets or slices from n=3 animals, Student t test p<0.001).

The granule fusion assay gives a quantitative measure of secretion but the low sample number of cells led us to directly measure insulin secretion using a bulk secretion assay. Here, we demonstrate that pancreatic slices, compared to isolated islets have, significantly increased insulin secretion at all concentrations of glucose (Fig 2 E)). Furthermore, we observed insulin secretion at low glucose (2.8 mM-5 mM) concentrations in slices that was not seen in isolated islets, and the overall EC_50_ for glucose dose dependence was different (Fig 2E, EC_50_ 10.2 mM for islets and 8.6 mM for slices). In control experiments, embedding isolated islets in agarose (the substrate used to embed slices) had no effect in insulin secretion (Supplemental Fig 2). Furthermore, there was no difference in measured proinsulin secretion in slices compared to isolated islets, indicating that insulin processing was unchanged (Supplemental Fig 2). Our values for insulin secretion in isolated islets are comparable with other reports (Rorsman and Ashcroft, 2018) and overall our data shows that slices are more sensitive to glucose and secrete more insulin than isolated islets.

A more detailed interrogation of glucose dependent control of insulin secretion segregates glucose action into two distinct routes: a trigger and an amplification pathway (Gembal et al., 1992; Henquin, 2009). The triggering pathway includes the steps from glucose uptake, closure of K_ATP_ channels and the activation of voltage dependent Ca^2+^ channels and the subsequent exocytosis of insulin granules (Rorsman and Ashcroft, 2018). Less is known about the amplification pathway which is characterised by a glucose dependent augmentation of insulin secretion (Gembal et al., 1992; Henquin, 2009) potentially by controlling granule transport and docking prior to fusion (Ferdaoussi et al., 2015). One approach to distinguish between the trigger and the amplification pathways uses diazoxide, a K_ATP_ channel opener, to clamp the β cell membrane potential negative prior to addition of glucose at different concentrations (Gembal et al., 1992). Glucose addition then does not cause insulin secretion, because of the presence of diazoxide, but secretion can be triggered by exposure to high potassium. Comparison of the responses at different glucose concentrations then defines glucose dependent amplification (Henquin, 2009).

In our experiments, because glucose-dependent secretion was greater in pancreatic slices at all glucose concentrations (Fig 2E), we were anticipating that amplification would be larger. Surprisingly, our results, showed the opposite and in fact glucose-dependent amplification was significantly larger in isolated islets compared with pancreatic slices (Fig 2F). This enhanced amplification suggests the overall decrease in glucose-dependent secretion in isolated islets, must be due to reduced glucose-dependent triggering.

These results demonstrate that at least one component of the enhanced secretion seen in pancreatic slices is due to β cell intrinsic differences. The mechanisms behind glucose-dependent amplification are not well understood and the increase in isolated islets, is therefore difficult to study. However, the steps in glucose dependent triggering are well understood and lead to Ca^2+^ responses. Given the enhancement in secretion in pancreatic slices we set out to characterise this glucose triggered Ca^2+^ signal in more detail.

### Glucose-dependent triggering: fast intracellular Ca^2+^ waves characterise responses in pancreatic slices

The final step in the glucose-dependent triggering pathway is the entry of Ca^2+^ through voltage-sensitive Ca^2+^ channels that open at each action potential (Rorsman and Ashcroft, 2018). We chose to study intracellular Ca^2+^ responses using the genetically encoded Ca^2+^ probe, GCAMP6s which was expressed in the β cells using knock-in InsCre mice (Thorens et al., 2015). The β cells were imaged using multiphoton microscopy and the responses across a range of glucose concentrations measured.

In slices (Fig 3A-D) and isolated islets (Fig 3E-H) we observed characteristic large responses to high glucose concentrations of 16.7 mM. When recording from different cells in the field of view we usually observed synchronous responses across many cells (Fig 3D, 3H) indicating that in both preparations the cells are functionally coupled through gap junctions (Benninger et al., 2011). In these recordings the Ca^2+^ responses from β cells within slices typically showed pulsatile activity even at 2.8 mM glucose (Fig 3D), which is consistent with the observations of insulin secretion at this low glucose concentration (Fig 2), and we also observed rapid pulsing of Ca^2+^ at the beginning of the high glucose induced responses in slices, consistent with enhanced excitability.

**Figure 3.**
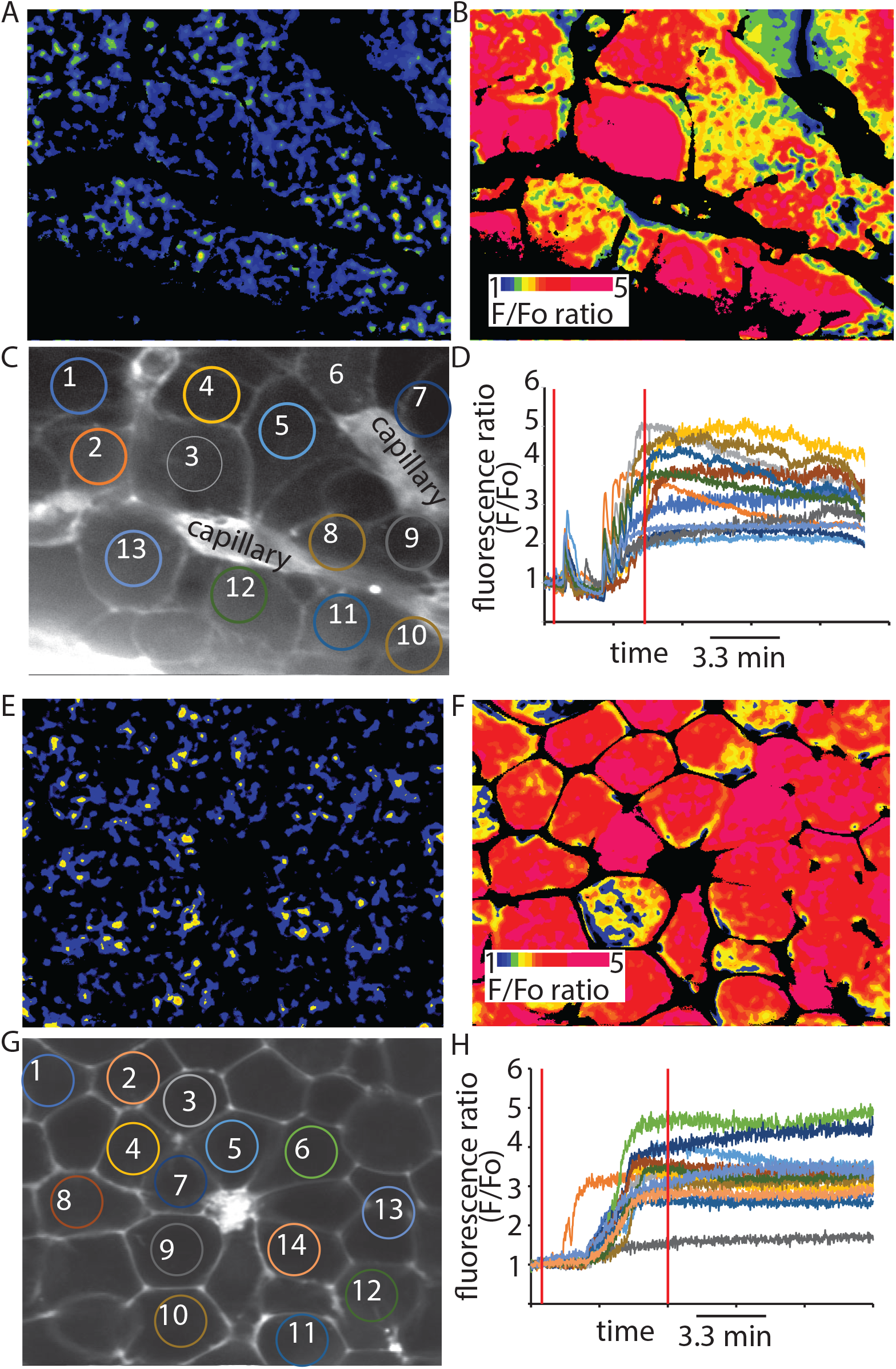
GCaMP6 recording show synchronous glucose induced Ca^2+^ responses in slices and isolated islets. GCaMP6 expressed in β cells shows rapid, synchronous Ca^2+^ responses in (A) slices) and (B) isolated islets in response to an increase of glucose concentration from 2.8 mM to 16.7 mM. In slices we consistently observed Ca^2+^ spikes at 2.8 mM and rapid Ca^2+^ jumps in the rising phase of the glucose-induced response.

We next recorded Ca^2+^ responses and determined the time when high glucose arrived at the cells by including a fluorescent probe in the high glucose solution (Supplemental Fig 4). Ca^2+^ responses in slices (Fig 4A,B) were apparently initiated almost simultaneously with the addition of 16.7 mM glucose indicating that these large responses are triggered by even small elevations in the concentration of glucose. In contrast, in isolated islets the Ca^2+^ responses occurred with a consistent delay after the addition of glucose (Fig 4C,D). Comparison of the parameters of the global Ca^2+^ responses to 16.7 mM glucose in slices with those in isolated islets shows the time to peak was significantly shorter in slices (Fig 4F).

**Figure 4.**
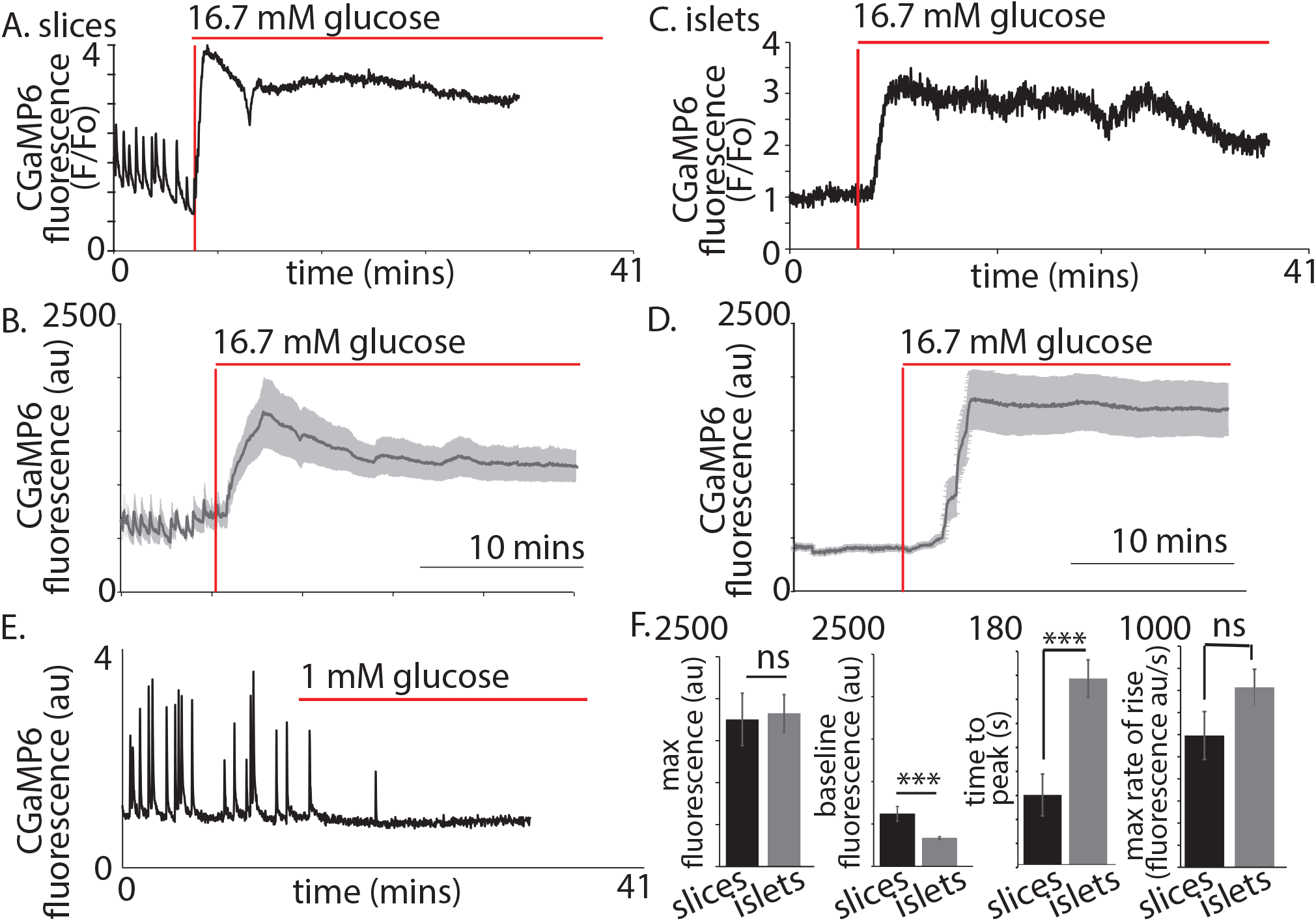
β cell Ca^2+^ responses in slices have short latencies to peak and higher glucose sensitivity compared to isolated islets. In slices, (A) single example or (B) averaged responses of Ca^2+^ measured by changes in GCaMP6 fluorescence in β cells within slices showed large, sustained responses to an increase of glucose from 2.8 mM to 16.7 mM. In slices we often observed fast Ca^2+^ spiking in β cells (5/7 slices) prior to the increase in glucose. In isolated islets the magnitude of (C) single responses, or the (D) average responses were similar to those in slices. (E) the Ca^2+^ spiking observed at 2.8 mM glucose from β cells within slices was lost when glucose was lowered to 1 mM. (F) the maximum fluorescence and overall maximum rate of rise (islets n=42 cells, 3 animals, in slices, n=26 cells, 3 animals, Student t test p=0.11). of the Ca^2+^ response was not different between slices and isolated islets. In contrast, the baseline fluorescence and the time to the peak Ca^2+^ response, from the addition of glucose, was significantly faster in slices vs islets (in isolated islets, n=43 cells, 6 islets, 3 animals and n=18 cells, in slices, n=18 cells, 4 slices, 3 animals, Student t test, p<0.001).

As before (Fig 3) we consistently observed pulsatile Ca^2+^ activity at 2.8 mM glucose (Fig 4A,B,E) which resulted in a significant elevation of the average “baseline” Ca^2+^ signal in slices compared with isolated islets (Fig 4F). These “baseline” Ca^2+^ pulses were glucose dependent and lowering glucose from 2.8 mM to 1 mM abolished all activity (Fig 4E).

We conclude that the Ca^2+^ responses observed at 2.8 mM glucose and the shorter latency to peak Ca^2+^ responses in the slices are consistent with the enhanced glucose-sensitive insulin secretion we observe (Fig 2) and confirm that it is the glucose dependent trigger that is enhanced in slices. This evidence indicates increased excitability in the Ca^2+^ pathway but does not suggest any mechanism that might underlie response. Furthermore, if insulin secretion was regulated by synaptic-like mechanisms then a key additional characteristic of synaptic control is that Ca^2+^ channels are locally regulated presynaptically to locally deliver Ca^2+^ to the sites of vesicle fusion. Interestingly, the preservation of the capillary bed in slices enabled us to determine the orientation of each β cell within the living slices and measure the Ca^2+^ responses in β cells adjoining the capillary. In these cells we often observed fast Ca^2+^ waves across the cell that originated at the capillary interface (Fig 5A-C). This indicates a spatial clustering of functional Ca^2+^ channels in the region adjoining the capillary.

**Figure 5.**
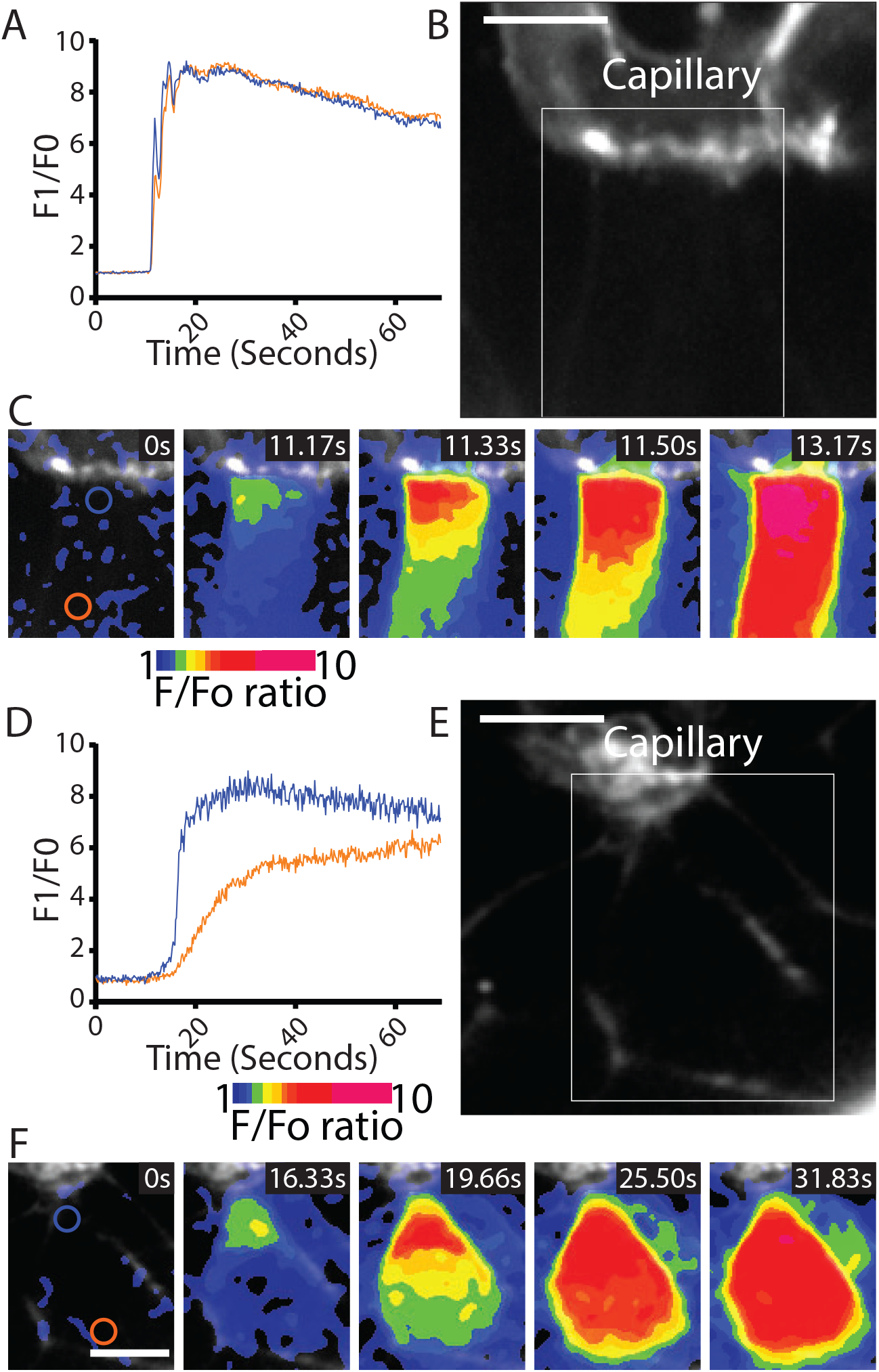
Fast Ca^2+^ waves originate at the capillary interface of β cells in slices. (A) β cells within the slices that adjoin the capillaries often showed Ca^2+^ responses that originated at the capillary interface and spread rapidly across the cell (apparent velocity 50.6+/-6.1 µm.S^-1^, mean+/-SEM, n=7 slices from 6 animals). (B) in isolated islets the capillaries were fragmented, and we rarely observed Ca^2+^ waves. The waves we did observe originated at the interface with capillary fragments and had a slow velocity. (1.8+/-0.2 µm.S^-1^, mean+/-SEM, n=3 islets from 3 animals, significantly slower compared with the velocity in slices, Student t test p<0.01).

In isolated islet preparations capillary structures were disrupted and observations of the Ca^2+^ responses in the adjoining cells showed that Ca^2+^ waves could be observed but these were rare (Fig 5D-F) and had a reduced wave velocity.

The observed Ca^2+^ waves, originating at the capillary interface, indicate mechanisms of locally increased Ca^2+^ channel activity in this region and are reminiscent of observations at the presynaptic domain. This regionally enhanced Ca^2+^ channel activity is likely to be controlled by protein complexes (Ohara-Imaizumi et al., 2019) and also by Ca^2+^-dependent feedback mechanisms that are intrinsic to channel control (Zühlke et al., 1999). Together these mechanisms could account for the increased excitability observed in the slices and the enhanced insulin secretion.

In neurones the presynaptic complex, including Ca^2+^ channels, is positioned through mechanisms that couple to the postsynaptic domain (Sudhof, 2012). In β cells there is no domain analogous to the postsynaptic region and therefore there must be alternative external environmental cues that position the presynaptic scaffold complex (Lammert and Thorn, 2020; Ohara-Imaizumi et al., 2019) and localise the control of the Ca^2+^ channel excitability that we have revealed. We next therefore tested the most likely of these cues, the extracellular matrix and the activation of the integrin/FAK pathway which we show is preserved in the slices (Fig 1).

### Integrin/focal adhesion control of glucose dependent Ca^2+^ signalling

FAK phosphorylation is enhanced by glucose stimulation and the small molecular inhibitor, Y15, significantly reduces phosphorylation (Rondas et al., 2012). In our experiments, pretreatment of slices with Y15 completely abolished the Ca^2+^ spikes observed at 2.8 mM glucose (Fig 6 A,B) and significantly reduced the responses to 16.7 mM glucose (Fig 6 C,D). Consistent with this inhibition, Y15 reduced glucose-induced insulin secretion in slices (Fig 6E) and interestingly had no effect on high potassium induced insulin secretion. We conclude that FAK is activated at the β cell capillary interface, the same region where Ca^2+^ signals originate, and that it selectively enhances glucose dependent Ca^2+^ channel excitability. To test this idea further we moved to an *in vitro* model.

**Figure 6.**
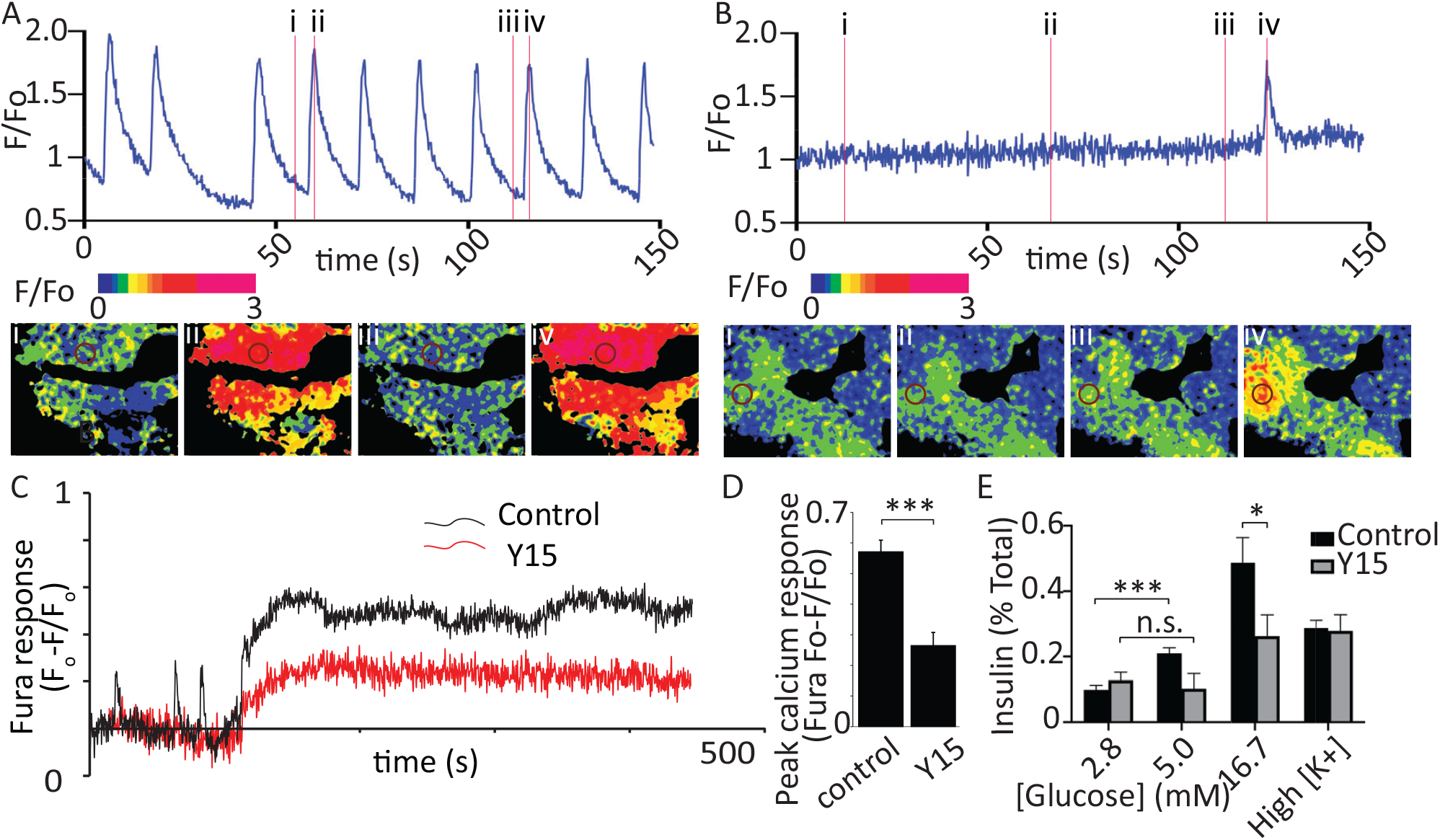
Focal adhesion kinase activation regulates glucose-induced Ca^2+^ responses. (A) As before, in the slice preparation, Ca^2+^ spikes were observed at 2.8 mM glucose, as measured with GCaMP6 fluorescence changes. (B) Pretreatment of slices with 2 µM Y15, an inhibitor of FAK, blocked these Ca^2+^ spikes (data from slices obtained from n=3 separate animals). To accurately measure the peak amplitude of Ca^2+^ responses we loaded cells with Fura-2, which has a lower Ca^2+^ affinity than GCaMP6 (Ca^2+^-induced fluorescence decreases are expressed as Fo-F/Fo to normalise for the initial fluorescence and to give positive deflections with increases in Ca^2+^). (C) Ca^2+^ responses to 16.7 glucose were robust in control and inhibited after pretreatment with Y15, with a significant reduction in peak amplitude (D, n=8 cells in slices from 3 separate animals, Student t test p<0.001). (E) insulin secretion, measured in slices, was blunted by pretreatment with Y15. In control, secretion was significantly increased (Student t test p<0.001, slices from n=3 mice) when stepping from 2.8 to 5 mM glucose and this increase was not seen in the presence of Y15. Responses to high potassium were not affected by the drug.

Culture of isolated β cells onto ECM coated coverslips is known to enhance overall insulin secretion (Parnaud et al., 2006) and through local integrin activation lead to targeting of insulin granule fusion to the interface of the cells with the coverslip (Gan et al., 2018). But how closely this replicates the polarisation seen in native β cells within slices has not been explored.

Here, we cultured isolated β cells on laminin coated coverslips and used immunofluorescence to determine if the structural response of the cells to contact with ECM mimicked that found in the native islet where the cells contact the ECM of the capillaries (eg Fig 1). The distribution of E-cadherin showed that cadherin interactions characterise cell-cell contacts (Fig 7A,B). Cells cultured on BSA (as an inert protein control) did not adhere well, they grew on top of each other and although phospho-FAK was apparent at the contact points of the cells with the coverslip it was sporadic and mainly on the outer edges of the cells (Fig 7A). In contrast, cells cultured on laminin grew as a monolayer with extensive punctate phospho-FAK staining at the footprint (Fig 7B). Immunostaining for the synaptic scaffold proteins liprin and ELKS (Fig 7C-H) showed significant enrichment at the coverslip interface when β cells were cultured on to laminin (Fig 7D-H) and not on BSA (Fig 7C-F); which is consistent with an integrin dependent mechanism of location both here and within slices (Fig 1).

**Figure 7.**
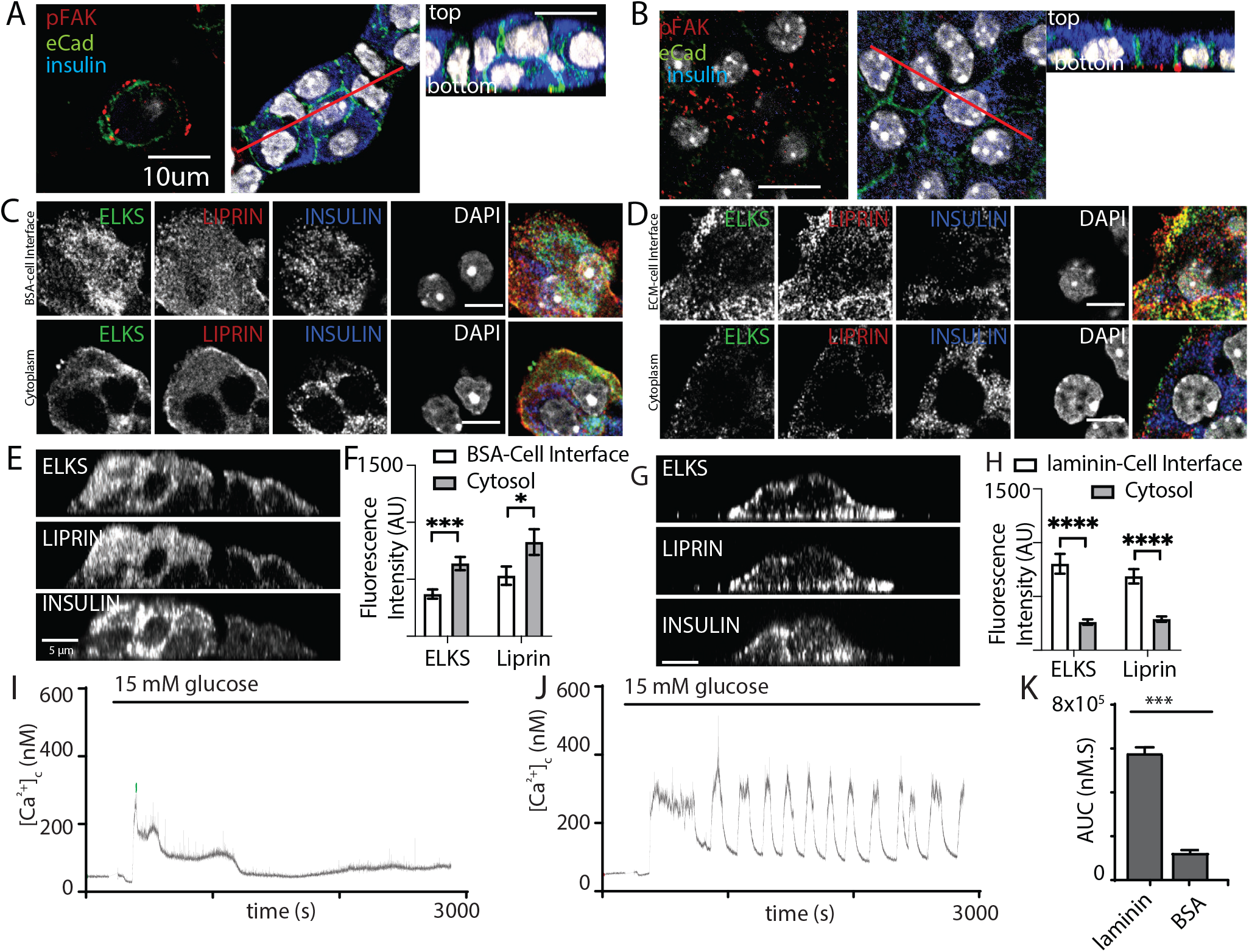
Integrin activation mediates β cell orientation and glucose dependent Ca^2+^ responses. (A) Immunofluorescence staining of phospho-FAK, E-cadherin and insulin showed that isolated β cells, cultured on BSA coated coverslips, were disorganised. Cells were multilayered and the phospho-FAK staining scattered at the edges of the footprint of the cells, also see orthogonal sections. (B) in contrast cells cultured on laminin coated coverslips showed extensive, punctate phospho-FAK located at the cell footprint (as shown in the orthogonal section) and organised E-cadherin staining at the cell junctions. (C-H) Immunofluorescence staining of isolated β cells (insulin; blue), grown on BSA (C) or laminin (D) coated coverslips showed enriched ELKS (green) and liprin (red) staining at the laminin-cell, but not BSA-cell interface, compared with the cytoplasm. This is illustrated in orthogonal sections (XZ) for cells cultured on BSA (E) or laminin (G). (F) Average fluorescence intensity of both ELKS (Student’s t test, p<0.001) and liprin (Student’s t test, p<0.05) were significantly lower at the BSA-cell interface compared with the cytosol (36 ROIs, n=6 cells from 3 animals). (H) In the cells cultured on laminin however, average fluorescence intensity of ELKS and liprin were significantly higher at the laminin-cell interface compared with the cytosol (Student’s t tests, p<0.001) (36 ROIs, n=6 cells from 3 animals). Using Fura-2 loaded, isolated β cells cultured on BSA, high glucose induced a modest, short-lasting response (I) that contrasted with the large response and sustained oscillations when the cells were cultured on laminin (J), with a significant reduction in area under the curve (AUC) of the response (K, student t test p<0.001, n=36 cells on laminin and n=21 cells on BSA). Scale bars 5µm.

This *in vitro* organisation of β cell structure therefore shares similarities with β cells in a slice including potentially a presynaptic-like domain. We therefore tested whether this would impact on the glucose dependent Ca^2+^ responses. β cells cultured on either BSA or on laminin showed glucose induced Ca^2+^ response (Fig 7I,J) but only cells on laminin showed robust long lasting Ca^2+^ oscillations and the overall AUC was significantly greater in the cells on laminin (Fig 7K).

This work is consistent with the observed effects of FAK inhibition on Ca^2+^ responses in slices (Fig 6) but we were concerned that there might be non-specific effects of the different culture conditions, for example the cells on BSA grow as three-dimensional clusters. To address this, we chose acute interventions applied to β cells cultured on laminin. In the first approach we pretreated the cultures with integrin β1 blocking antibodies and, consistent with the data in Fig 7 we saw both a disruption in the localisation of liprin at the coverslip interface and an inhibition of the glucose induced Ca^2+^ responses (Supplemental Fig 7).

In the second approach we used the FAK inhibitor Y15 applied to β cells cultured on laminin coated coverslips (Fig 8). In the presence of Y15 the glucose induced Ca^2+^ response was significantly reduced (Fig 8A-C) and glucose induced secretion, but not high potassium, was also inhibited in a reversible manner (Fig 8D); both consistent with the actions of Y15 in the slices (Fig 6). Immunofluorescence studies showed that the distribution of liprin and ELKS were disrupted by Y15 (Fig 8E-J), consistent with the data showing the importance of the integrin/FAK pathway in their positioning.

**Figure 8.**
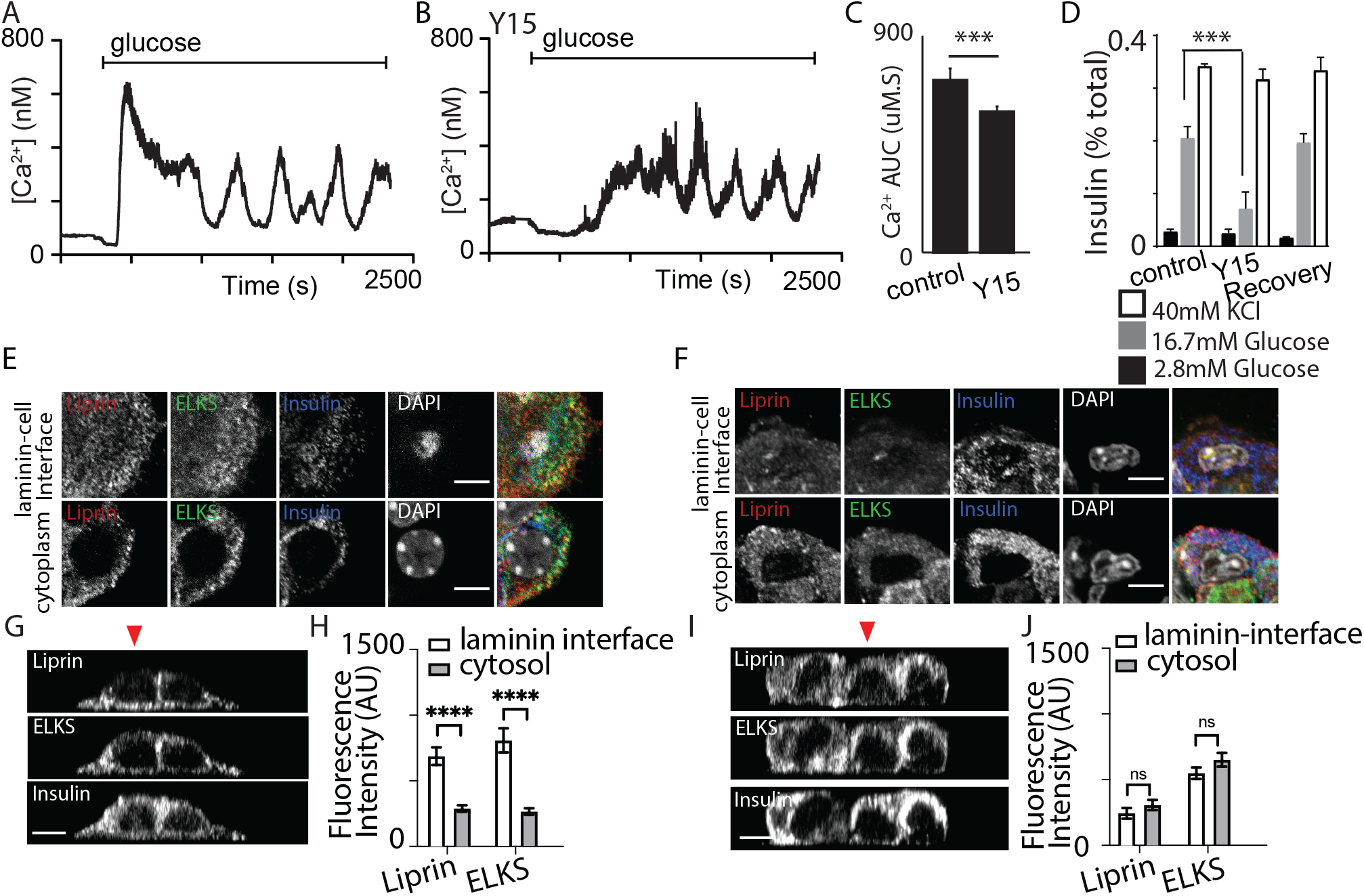
FAK regulates both Ca^2+^ responses and positioning of presynaptic scaffold proteins. Isolated β cells were cultured on laminin coated coverslips and FAK was inhibited by pretreatment with 2 µM Y15. In Fura-2 loaded cells we observed the typical robust response to high glucose followed by sustained oscillations in control (A). A smaller, delayed response was observed in the presence of Y15 (B) with a significant reduction in AUC (C, using regions of interest from n=218 cells in DMSO and 208 cells in Y15, from 3 mice, Student’s t test p<0.001). (D) consistent with this action of Y15 we observed a reversible reduction in glucose induced insulin secretion in the presence of Y15 (n=3 animals in each condition, Student’s t test p<0.001). However, no significant difference in insulin secretion was observed following potassium stimulation between cells incubated with Y15 compared with DMSO control (n = 3 animals, Student’s t test p=0.25). (E,G,H)) as before, immunostaining showed enrichment of liprin and ELKS at the laminin-cell interface which was blocked after pretreatment with Y15 (F,I,J, ELKS Student’s t test, p = 0.15 and liprin, Student’s t test, p = 0.28, 36 ROIs, n=6 cells from 3 animals). Scale bars 5µm.

Taken together our data provides strong evidence that the integrin/FAK pathway is critical both for the local enrichment of synaptic scaffold proteins in β cells and for locally enhanced excitability of the Ca^2+^ channels.

## Discussion

Our interrogation of β cell structure and function in pancreatic slices shows precise sub-cellular organisation, targeting of granule fusion to the capillary interface and enhanced insulin secretion that points to a robust glucose dependent trigger. We observe Ca^2+^ spikes at low glucose and short-latency responses to high glucose showing enhanced sensitivity of the cells to glucose in slices compared to isolated islets. Using a range of interventions, we show a glucose dependent integrin/FAK pathway locally enhances the Ca^2+^ response and positions the presynaptic scaffold proteins, ELKS and liprin. This work demonstrates that the FAK pathway intersects with the final stages of the glucose dependent control of secretion and has important implications for our understanding of the stimulus secretion cascade in β cells and treatments for diabetes.

### FAK and the control of insulin secretion

We are not the first to identify a role for FAK in the control of secretion. Halban’s group showed in mouse β cells that FAK phosphorylation was increased by glucose stimulation (Rondas et al., 2011) and block of integrins or FAK inhibited insulin secretion from the MIN6 cell line with evidence that it affected F-actin remodelling (Rondas et al., 2012). In a mouse study, knockout of FAK caused hyperglycemia and using isolated islets they showed a reduced insulin secretion but no effect on Ca^2+^ responses (Cai et al., 2012). However, our work now shows that we must be careful in interpreting data from isolated islets; the dramatic reduction in phospho-FAK compared to slices (Fig 1) means the integrin/FAK pathway is compromised, even in controls. Interestingly, Halban’s approach cultured the β cells onto dishes coated with extracellular matrix (Rondas et al., 2011) which, since we now demonstrate is an excellent model that recapitulates FAK activation, β cell organisation, Ca^2+^ signals and secretory responses, is a much better approach to explore this pathway.

These previous studies did not explore the sub-cellular actions of the integrin/FAK pathway and, although they imply an action on F-actin, the mechanism was not explored. In contrast, we show direct evidence that FAK is a master regulator of two processes in the latter stages of glucose dependent control of insulin secretion where it controls the positioning of presynaptic scaffold proteins and controls the Ca^2+^ signal.

### Evidence that the integrin/FAK pathway regulates synaptic-like mechanisms to control insulin secretion

In neurones the key steps from opening of voltage gated Ca^2+^ channels to the exocytic fusion of vesicles are tightly spatially regulated by presynaptic complexes that are also the site for modulation of responses (Sudhof, 2012). In β cells, closely analogous steps use glucose dependent Ca^2+^ signals to induce insulin granule fusion, furthermore, presynaptic scaffold proteins are present (Low et al., 2014; Ohara-Imaizumi et al., 2005) and function to control insulin secretion (Fujimoto et al., 2002; Ohara-Imaizumi et al., 2005; Shibasaki et al., 2004). However, whether these scaffold proteins exist as a complex that regulate insulin secretion in a manner analogous to synaptic control is not clear.

Here we provide evidence that aspects of the control of insulin secretion in β cells are similar to presynaptic mechanisms. We show that presynaptic scaffold proteins, insulin granule fusion and the control of Ca^2+^ channels all occur locally where the β cells contact ECM. Furthermore, activation of the integrin/FAK pathway is critical for each one of these factors, either in positioning of granule fusion as we have previously shown (Gan et al., 2018) or, as we now show in the positioning of the scaffold proteins and regulation of the Ca^2+^ response.

In terms of spatial constraints, liprin, ELKS and other presynaptic scaffold proteins are all enriched at the capillary interface (Fig 1 (Low et al., 2014)) and when this complex is preserved, as we now show in slices, there is a very tight focus of insulin granule fusion to this region (Fig 2). This is consistent with a synaptic-like mechanism. The various roles of liprin in neurones are still being uncovered but through protein-protein interactions it nucleates the formation of the presynaptic complex including proteins such as RIM which in turn tether granules (Sudhof, 2012; Wei et al., 2011). Future work will be required to identify if liprin plays a similar role in β cells.

One point of distinction in the β cell compared to neurones is that there is no equivalent to a post-synaptic domain. In neurones the pre and post synaptic domains are aligned by transmembrane proteins that span the synaptic cleft, such as neurexins (Sudhof, 2008). Indeed, neurexins do exist in β cells (Mosedale et al., 2012) but our work now suggests that the integrin/focal adhesion pathway is a more likely candidate controlling the positioning of the presynaptic complex and we directly show it controls the positioning of both ELKS and liprin. The question arises as to how this occurs and although there is evidence that liprins do interact with focal adhesions (Astro et al., 2016) this has not been explored in β cells.

In terms of control of the Ca^2+^ response, our new evidence indicates that synaptic-like mechanisms play a role. The Ca^2+^ responses we observe are a spatial and temporal integration of discrete bursts of Ca^2+^ entry at each action potential (Rorsman and Ashcroft, 2018). Our data show that the maximal global rate of rise of the GCaMP measured Ca^2+^ response is similar between the slices and isolated islets (Fig 4F). This suggests that the number of active Ca^2+^ channels in the β cells in both preparations is similar and is therefore consistent with the long-standing observations of robust Ca^2+^ responses in isolated islets. What is different in the slices is that we observe rapid local increases in Ca^2+^ and waves at the capillary interface, which must reflect local clustering of active channels – a central characteristic of neuronal synapses.

How do we explain the enhanced sensitivity to glucose of the Ca^2+^ responses in slices? Specifically, we might expect mechanisms that act on the voltage sensitivity of the Ca^2+^ channels, so they respond at more negative membrane potentials, or that the Ca^2+^ channels open longer and increase Ca^2+^ influx. Our data provides evidence for two possible factors that are shaping the Ca^2+^ responses in slices. Firstly, the clustering of active Ca^2+^ channels at the capillary interface will affect Ca^2+^ channel behaviour. In the mouse the predominant Ca^2+^ channel is Cav1.2 (Schulla et al., 2003) which is positively and negatively regulated by cytosolic Ca^2+^ (Zühlke et al., 1999). As has been shown in many other systems, the entry of Ca^2+^ through each channel influences its own activity and the activity of immediately surrounding channels which makes channel clustering a critical factor in controlling channel opening (Stanley, 1997). Secondly, the localised activation of focal adhesions (Fig 1), targets Ca^2+^ channels. We show that culture of cells on BSA, inhibition of FAK and integrin β1 blockade all reduce the Ca^2+^ response to glucose. This is the first report of a link between integrins and Ca^2+^ response in β cells, which could be mediated through signal cascades elicited by focal adhesion activation, as has been shown in smooth muscle cells (Hu et al., 1998) or it could be secondary to an integrin/FAK mediated positioning of synaptic scaffold proteins. For the latter, we have shown integrin activation positions liprin and ELKS (Fig 8) and in turn ELKS may position the Ca^2+^ channels (Ohara-Imaizumi et al., 2019).

### Enhanced sensitivity to glucose in slices

Our finding of enhanced sensitivity to glucose in the pancreatic slices is a significant advance in the field. We observe repetitive Ca^2+^ spikes at 2.8 mM glucose that are lost when glucose is lowered to 1 mM and are not seen in isolated islets. In parallel, insulin secretion is observed from slices at 2.8 mM and decreases when glucose is lowered. This enhanced glucose sensitivity is likely to be driven by the intrinsic factors within the β cells we have identified. These factors include the identification of fast Ca^2+^ waves that originate at the capillary interface, the short latency to peak Ca^2+^ responses and the close coupling between the Ca^2+^ signals and sites of insulin granule exocytosis. We cannot rule out that other factors, present in pancreatic slices, may influence glucose sensitivity. One possible factor is the gap junction coupling of the cells, where, at low glucose concentrations, a majority of non-responsive cells are thought to supress the activity of individual particularly excitable cells (Benninger et al., 2011). However, this does not seem a likely explanation for our findings because we observe strong coordination of Ca^2+^ responses, indicative of cell-to-cell coupling, in both slices and isolated islets. Another obvious factor, that might differ in the preparations, are α cells where glucagon secretion can stimulate insulin release (Moede et al., 2020). However, this seems unlikely because lowering glucose from 2.8 to 1 mM would stimulate glucagon secretion and in the β cells we observe the opposite; a reduced insulin secretion and a reduced Ca^2+^ response.

In a broader physiological context, it might seem unlikely that the responses we observe to low glucose concentrations are real. The “set point” for mouse blood glucose is ∼7 mM (Rodriguez-Diaz et al., 2018) and the consensus from other studies, mostly using isolated islets, is that insulin secretion has an EC_50_ for glucose of ∼8 mM (Hedeskov, 1980). Furthermore, the K_m_ for the GLUT 2 transporter is 11 mM and the EC_50_ for mouse glucokinase is 8 mM (Rorsman and Ashcroft, 2018). However, there is precedent that β cells can respond to much lower glucose concentrations. Henquin’s lab showed a dose-dependence of the amplifying pathway from 1-6mM (Gembal et al., 1992) and extensive early work identified subpopulations of isolated β cells that are very sensitive to glucose and released maximal insulin at 8.3 mM glucose (Van Schravendijk et al., 1992), similar to our findings (Fig 2). Given the excellent preservation of cell structure within the slice, our results likely reflect optimal behaviour of β cells. Furthermore, *in vivo* glucose control reflects a balance of hormones and, at low glucose concentrations glucagon might be the dominant hormone but altered insulin secretion could also play a role (Rodriguez-Diaz et al., 2018).

### Broader significance

Our work has important implications for understanding and treating diabetes. For type 2 diabetes, past work has indicated an impact of lipotoxicity on Ca^2+^ channel organisation (Hoppa et al., 2009) and the disease on Ca^2+^ clustering (Gandasi et al., 2017) which, in the new context given by our work, would take place at the capillary interface. We also know that both the capillary structure (Brissova et al., 2015) and the extracellular matrix composition (Hayden et al., 2005) are altered in disease. In the light of our work, it is likely that this will affect the β cell responses through a disruption of the integrin/FAK pathway we describe. Given that sulfonylureas can improve insulin secretion in T2D (Rorsman and Ashcroft, 2018) we already know that enhancement of glucose-dependent triggering is beneficial. Our new work suggests that widening the scope of our interest to include each element of the triggering pathway would be fruitful and that specifically intervening with the primary mechanisms that spatially organise the β cells could be disease modifying.

For type 1 diabetes, exciting advances are leading to the development of stem cell based β cell replacements (Melton, 2021). Most approaches generate spheroids of cells that we have recently shown do not contain organised extracellular matrix (Singh et al., 2021) and, as a result, the β-like cells within the spheroids are not polarised (Singh et al., 2021). Our work now suggests that amplification will be the dominant pathway underpinning glucose-dependent insulin secretion in these spheroids and that these cells will lack a drive from the integrin/FAK pathway. Because the triggering and amplification pathways are distinct our work indicates that a selective focus on enhancement of triggering may be broadly beneficial. This could include imposing polarity to the β-like cells, which we have shown does enhance secretion (Singh et al., 2021), but it could also include genetic manipulation to upregulate components of the triggering pathway or the use of drugs, like sulfonylureas, to increase the sensitivity of this pathway.

## Materials and Methods

### Animal husbandry

Male C57BL6 and GCAMP-InsCre mice were ordered from Animal Bioresources and housed at the Charles Perkins Centre facility in a specific pathogen-free environment, at 22°C with 12-hour light cycles. All mice were fed a standard chow diet (7% simple sugars, 3% fat, 50% polysaccharide, 15% protein (w/w), energy 3.5 kcal/g). Mice (8-12 weeks old) were humanely killed according to local animal ethics procedures (approved by the University of Sydney Ethics Committee).

### Glucose-stimulated insulin secretion (GSIS) and HTRF insulin assay

GSIS media was Krebs-Ringer Bicarbonate solution of pH7.4 buffered with HEPES (KRBH), plus 2.8mM glucose (basal) or 16.7mM glucose (stimulation). Depolarisation media was a modified KRBH with reduced NaCl (100mM) and high potassium (40mM KCl). Where applied, diazoxide (Sigma) was used at a concentration of 250 uM. All media and cells were kept at 37°C for the duration of the assay. Tissues were washed in warm basal media two times and then placed in fresh basal media for one hour. The basal media was washed out an additional time and then tissues were incubated for 30 minutes in fresh basal media. Tissues were collected at the end of the assay into ice-cold lysis buffer (1% NP-40, 300mM NaCl, 50mM Tris-HCl pH 7.4, protease inhibitors) and sonicated. Supernatants and lysates were stored at -30°C prior to HTRF assay (Mouse ultrasensitive, Cisbio).

### Islet preparation

Isolated mouse islets were prepared according to a standard method that utilizes collagenase enzymes for digestion and separation from exocrine pancreatic tissue (Li et al 2009). In brief, a Liberase (TL Research grade, Roche) solution was prepared in un-supplemented RPMI-1640 (Gibco) media at a concentration of 0.5 U/mL. Pancreases were distended by injection of 2 mL of ice cold Liberase solution via the pancreatic duct, dissected and placed into sterile tubes in a 37°C shaking water bath for 15 minutes. Isolated islets were separated from the cell debris using a Histopaque (Sigma) density gradient. Isolated islets were maintained (37°C, 95/5% air/CO_2_) in RPMI-1640 culture medium (Sigma-Aldrich), 10.7 mM glucose, supplemented with 10% FBS (Gibco, Victoria, Australia), and 100 U/ml penicillin/0.1mg/ml Streptomycin (Invitrogen, Victoria, Australia).

### Islet slices

Sectioning of unfixed pancreatic tissue was performed as described by Huang et al (Gan et al., 2017; Huang et al., 2011). Pancreatic sections (200 μm thick) were cut and incubated overnight in RPMI-1640 supplemented with penicillin-streptomycin, 10% FBS, and 100 ug/mL soybean trypsin inhibitor (Sapphire Bioscience).

### Tissue fixation and immunofluorescent staining

Tissues were fixed with 4% paraformaldehyde (Sigma-Aldrich) in PBS 15 minutes at 20°C. Samples were stored in PBS at 4°C prior to immunofluorescent staining. Immunofluorescence was performed as described by Meneghel-Rozzo et al. (Meneghel-Rozzo et al., 2004). Tissues were incubated in blocking buffer (3% BSA, 3% donkey serum, 0.3% Triton X-100) for a minimum of one hour at room temperature followed by primary antibody incubation at 4°C overnight. Sections were washed in PBS (4 changes over 30 minutes) and secondary antibodies (in block buffer) were added for 4 hours (whole islets and slices) or 45 min (cells) at 20°C. After washing in PBS, tissues were mounted in Prolong Diamond anti-fade reagent (Invitrogen).

### Imaging

Confocal imaging was performed on a Nikon C2 microscope using a 63x oil immersion objective or on a Leica SP8 microscope with a 100X oil immersion objective. Live-cell imaging was possible on a two-photon microscope constructed in-house using Olympus microscope components. Two-photon imaging was performed at 37°C. Images were analysed using ImageJ and MetaMorph software. A 3D circumference linescan analysis (for example in Fig 1K, L) used linescans around the cell circumference at each Z section. The fluorescence intensity along each circumference linescan was then plotted out as intensity plots to produce the 3D heatmaps. The heatmap was produced in Excel by assigning pseudocolors to fluorescence intensity. Quantitation of protein area (Fig 1C, D) was calculated by converting single channels to binary images using a threshold that eliminated background (estimated as the average signal in the area of the nucleus) and was normalised to total cell area.

### Islet slices and Fura-2 measurement

Slices were removed from overnight culture media and incubated in 6 well plates containing 1mL KRBH 11mM glucose with 6 µm Fura 2-AM, 2 slices per well on a rocking platform at room temperature for 1 hour. After incubation, slices were placed back in culture media and washed for up to 6 hours in an incubator set to 37C and 5% CO2. Slices were removed for experimentation as needed and imaged after a pre-basal period of 1 hour in KRBH 2.8 mM glucose with or without the presence of 2 µm Y15 in an incubator. After pre-basal, period single slices were removed and placed in a pre-heated imaging chamber at 37°C with 1 ml KRBH 2.8 mM glucose. Slices were stimulated by adding glucose solution to a final concentration of 16.7mM and imaged with an excitation laser tuned to 810 nm on a 2-photon microscope and emitted light collected between 470-520 nm.

### Antibodies

Primary antibodies used for this study were: anti-insulin (Dako Cytomation, A0564), anti-beta1 laminin (Thermo Scientific MA5-14657), anti-integrin beta 1 (BD Biosciences 555002), anti-talin (Sigma-Aldrich T3287), anti-phosphorylated FAK (Cell Signalling Tech 8556S), anti-liprin alpha1 (Proteintech 14175-1-AP), anti-ELKS (Sigma, E4531). All primary antibodies were diluted 1/200. Secondary antibodies were highly cross absorbed donkey or goat antibodies (Invitrogen) labelled with Alexa 488, Alexa 546, Alexa 594, or Alexa 647. All were used at a 1/200 dilution. DAPI (Sigma, 100 ng/ml final concentration) was added during the secondary antibody incubation.

| Target | Species | Manufacturer / Catalogue number |
| --- | --- | --- |
| Insulin | Guinea pig | DAKO, A0564 |
| Laminin-beta1 | Rat | Invitrogen, MA5-14657 |
| Integrin-beta1 | Hamster | BD Biosciences, 555002 |
| Liprin-alpha1 | Rabbit | Proteintech, 14175-1-AP |
| PAR3 | Rabbit | Millipore, 07-330 |
| E-cadherin | Mouse | BD Biosciences, 610181 |
| Phospho-FAK (Y397) | Rabbit | CST, 8556S |

### Islet cell seeding procedure

Single cell suspensions were prepared by digesting isolated islets with TrypLE express enzyme (Gibco). Culture medium was RPMI-1640 supplemented with 10% FBS, and 100 U/ml penicillin/0.1mg/ml Streptomycin. Cells were cultured in standard incubator conditions (37°C, 10% CO2, humidity 20%).

In most experiments (Fig 7) we simply used plain coverslips but in the insulin secretion assays (Fig 8D), to create a more stable covalent attachment of basement membrane proteins to the surface of the glass coverslips we coated the coverslips with a thin layer (approximately 10-20 nm thick) of plasma activated coating. The plasma treatment was conducted using a radio frequency (RF) power supply (Eni OEM-6) powered at 13.56 MHz and equipped with a matching box. Plasma ions were accelerated by the application of negative bias pulses from RUP6 pulse generator (GBS Elektronik GmbH, Dresden, Germany) for 20 μs duration at a frequency of 3000 Hz to the stainless-steel sample holder. Glass coverslips were first activated in argon plasma powered at 75 W under a 500 V negative bias for 10 minutes at 80 mTorr. After that, a gas flow consisting of acetylene (1 sccm), nitrogen (3 sccm) and argon (13 sccm) was introduced into the chamber for 10-minute plasma deposition. During this step, plasma was generated with 50 W RF power at a pressure of 110 mTorr while positive ions were deposited on glass cover slips under a negative bias of 500 V. After the plasma treatment, samples were kept in a petri dish in ambient condition until use.

PIII-treated coverslips or plain coverslips were coated with Laminin 511 (BioLamina) 5ug/ml or bovine-serum albumin (Sigma) 1 mg/mL overnight at 4°C. After coating coverslips were rinsed in in PBS and then the cells were seeded.

### Statistical Analyses

All numerical data are presented as mean +/- standard error of the mean. Statistical analysis was performed using Microsoft Excel and GraphPad Prism. Data sets with two groups were subjected Student’s t-test, significance is indicated as follows: * p<0.05, ** p<0.01, *** p<0.001.

## Acknowledgements

We acknowledge project funding obtained from the National Health and Medical Research Council (APP1128273, to PT), The University of Sydney Strategic Research Excellence Initiative (SREI to PT and MB), Diabetes Australia (DART grant to PT) and Australian Research Council (FL190100216 to MB). Imaging was performed in the Centre for Microscopy and Microanalysis at the University of Sydney.

## Author Contributions

NH, DJ, KD and PT conceptualised the aims and designed experiments. NH, WJG, DJ, KD, KK, JT CT all performed the experiments and analysis. CT and MB provided plasma treated materials. PT instigated and supervised all aspects of the project. PT initiated the writing of the manuscript and all authors participated.

---

**Supplemental Fig 1.**
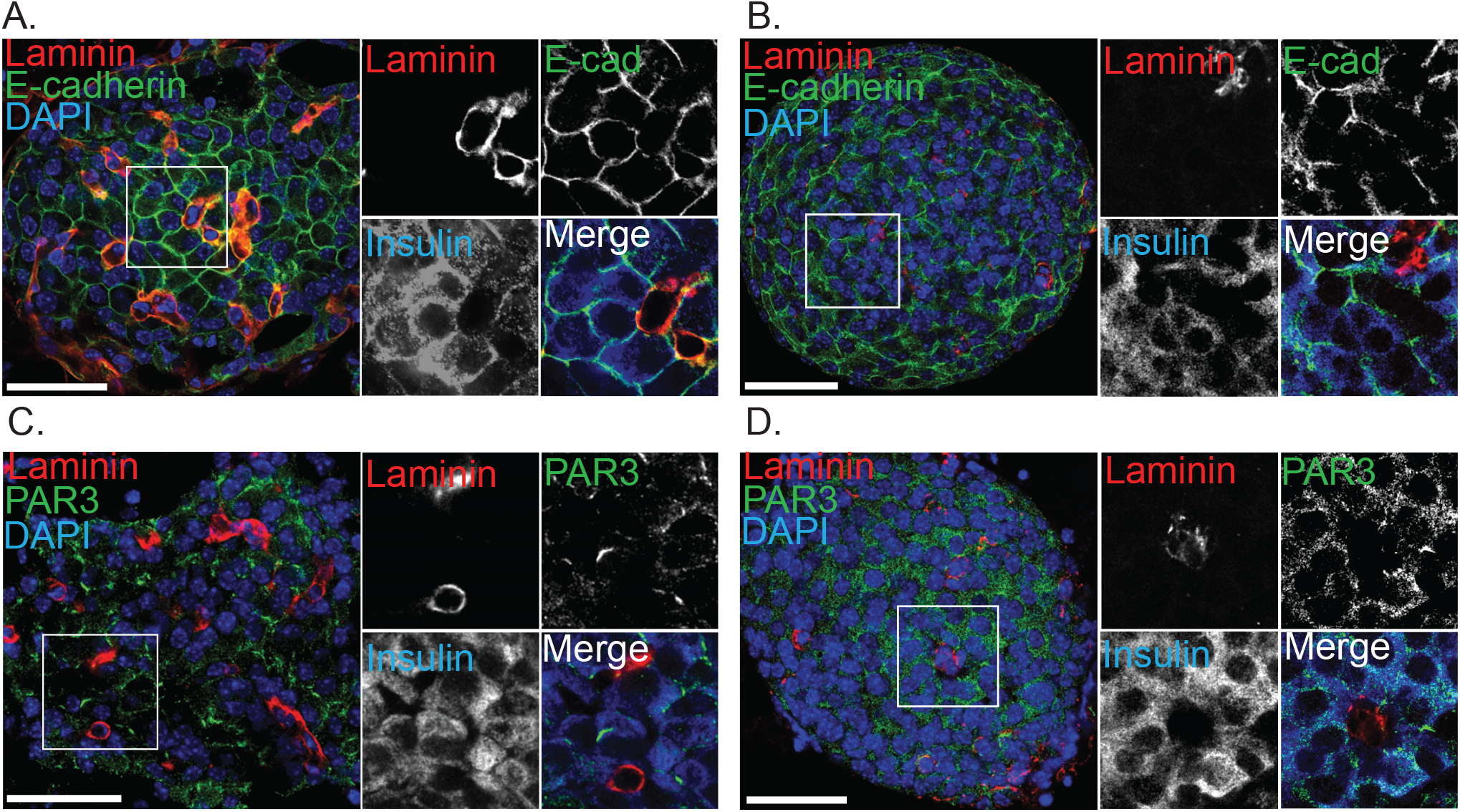
Immunostaining for E-cadherin and PAR3 in slices and isolated islets. Immunostaining for E-cadherin and laminin in (A) slices and (B) isolated islets shows enrichment at cell-cell interfaces in both preparations. Immunostaining for PAR3, the apical polarity determinant shows in (C) slices an enrichment in discrete regions away from the capillaries (stained with laminin) that is lost in (D) isolated islets. Scale bar 50 µm.

**Supplemental Fig 2.**
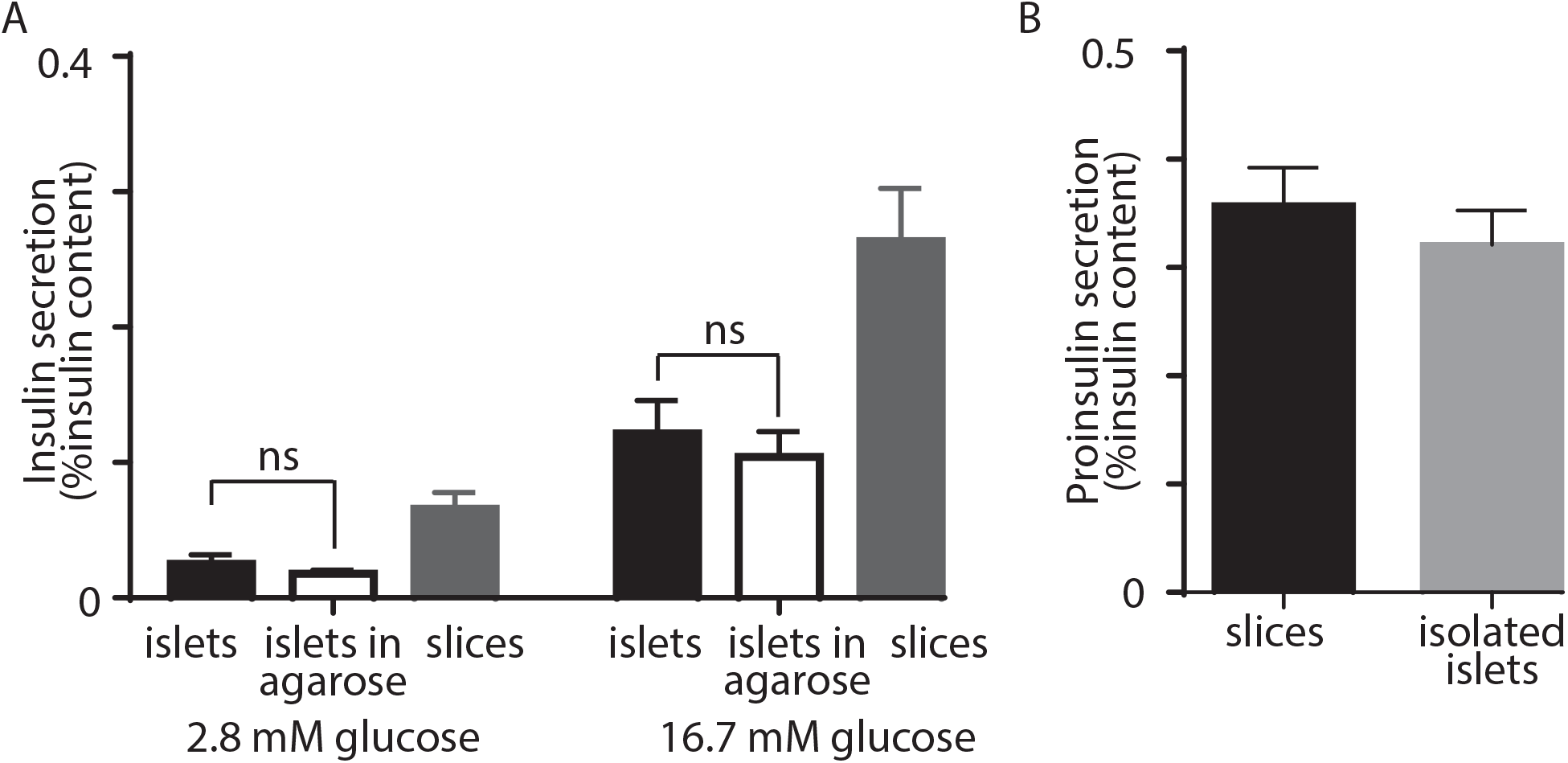
Insulin secretion in pancreatic slices and isolated islets. (A) isolated islets, embedded in agarose showed no difference (Student t test) from isolated islets alone in terms of basal and glucose stimulated insulin secretion. (B) measures of proinsulin secretion showed no difference in glucose stimulated slices versus isolated islets.

**Supplemental Fig 4.**
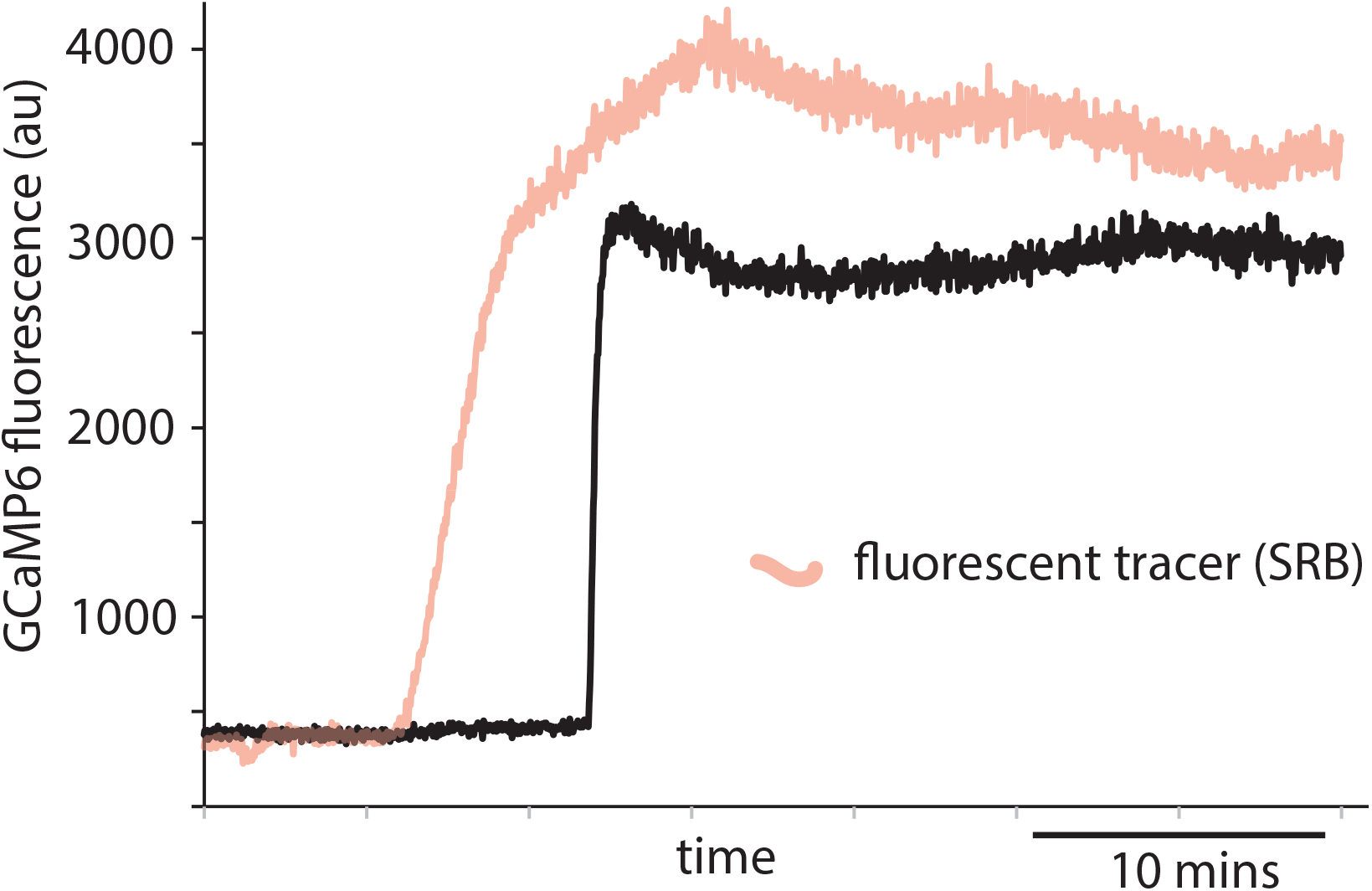
Example record showing use of fluorescent tracer to indicate addition of high glucose. The record shows a single cell Ca^2+^ response recorded from an isolated islet. The glucose concentration was changed from 2.8 mM to 16.7 mM where the high glucose solution contained sulforhodamine B as a fluorescent tracer which was recorded from a region of interest close to the responding cell. The time point where the SRB fluorescence increased identifies when the glucose concentration started to increase and was used to calculate the latency to the peak Ca^2+^ response.

**Supplemental Fig 7.**
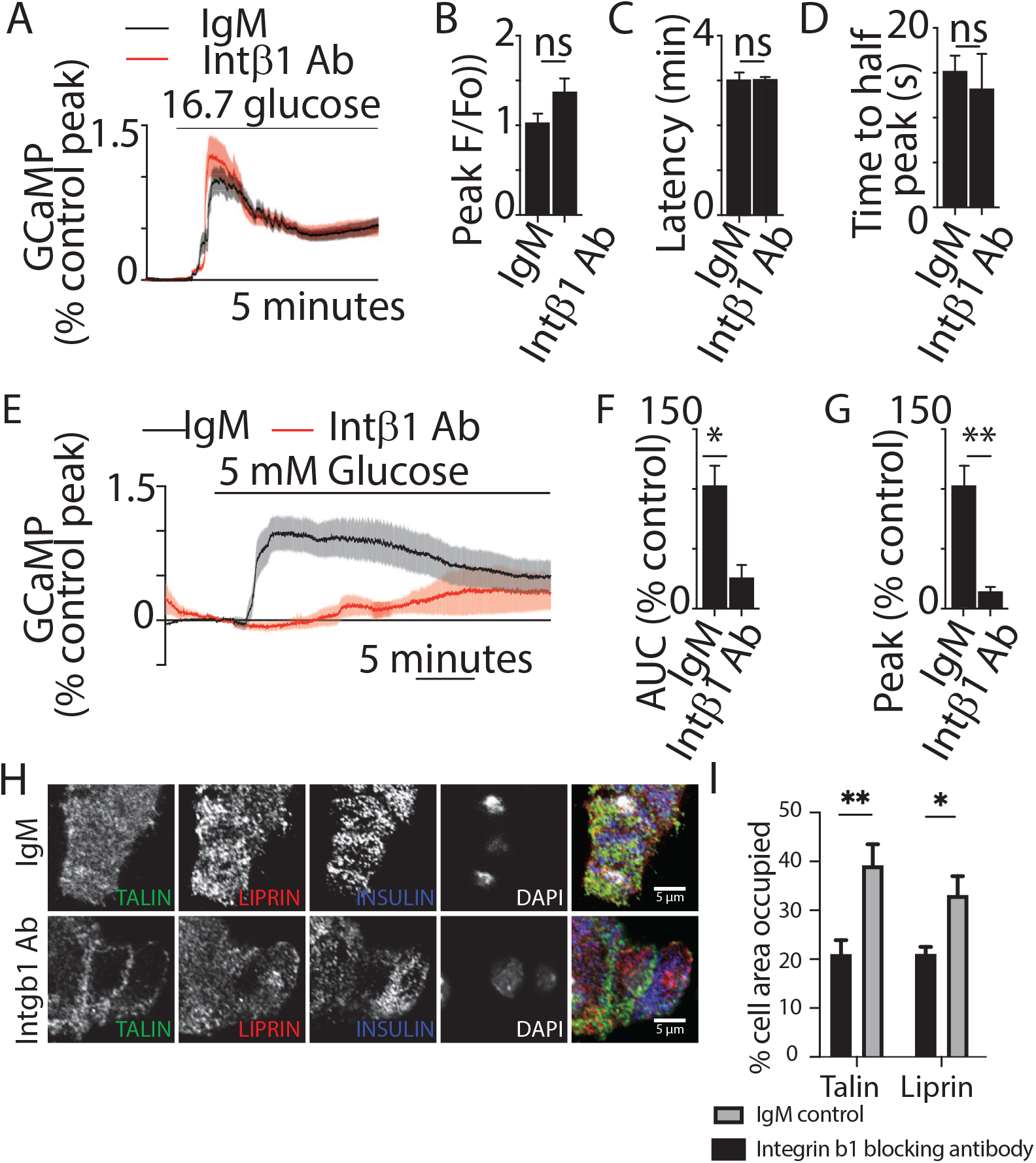
Blockade of integrin activation disrupts β cell structure. Glucose-induced Ca^2+^ responses were recorded using live-cell two-photon microscopy in β cells expressing the Ca^2+^ indicator GCaMP. GCaMP fluorescence was recorded over 30 minutes following high glucose (16.7mM) stimulation in isolated β cells incubated with integrin-β1 function blocking antibody compared with IgM control (for each condition, n = 47-49 cells from 3 animals). (A-D) No significant differences in GCaMP peak amplitude (Student’s t test, p = 0.09), latency (time between glucose addition and calcium response) (Student’s t test, p = 0.93), and time to half peak (Student’s t test, p = 0.66), were observed. (E-G) Representative Ca^2+^ traces within a single β cell, in the presence of IgM control or integrin-β1 function blocking antibody, in response to 5mM glucose (for each condition, n= 23-43 cells across 3 animals). (E) Average Ca^2+^ traces showed a robust response in the IgM condition, but a smaller and slower response in the Intβ1 block condition. (F,G) Incubation in Intβ1 blocking antibody decreased average AUC (Student’s t test, p < 0.05) and GCaMP peak amplitude (Student’s t test, p <0.01). Scale bars 5 µm. (H) immunostaining of β cells cultured on to laminin coated coverslips, showed that the positioning of liprin and the focal adhesion protein talin, that are normally enriched at the footprint, as shown by percentage of area occupied (I), are lost after preincubation with integrin β1 blocking antibody (Talin Student t test, p<0.01 and Liprin p<0.05, 6 cell clusters from 3 mice). Scale bar 5 µm.

